# A Near Telomere-to-Telomere Reference Genome for Drakensberger Cattle Achieved Through Trio-Binned Long-Read Sequencing

**DOI:** 10.1101/2025.11.19.689200

**Authors:** Ntanganedzeni Mapholi, Thendo Tshilate, Rae Smith, Nompilo Hlongwane, Lucky Tendani Nesengani

## Abstract

Drakensberger cattle is indigenous to South Africa and represent a unique genetic resource which is known for its quality beef production and adapted to the harsh climatic conditions. Despite its significance within the beef industry, no high-quality chromosomal level genome has been reported for this breed, limiting genomic selection and conservation efforts. Although there is a bovine genome reference, we need a breed specific reference to identify breed-unique alleles and structural variants that might not be explained by a distant reference. To address this, we generated a haplotype resolved assembly for Drakensberger cattle using a trio-binning where we employed PacBio HiFi reads for sire and dam, a combination of PacBio HiFi reads, Oxford Nanopore Technologies reads and Omni-C reads for the offspring. The assembled diploid genome size is 2.89 Gb with a scaffold N50 of 111 Mb and contig N50 of 61 Mb. The consensus accuracy was exceptionally high (QV = 70.23) and a genome completeness of 97.5 %, as analysed by the Benchmarking Universal Single-Copy Orthologs (BUSCO). The k-mer profiling suggested one of the strong haplotype separation for livestock with the paternity assembly containing 97.1% and maternity with 99.0%, combined diploid genome recovered 99.46% of all the solid read k-mers. We identified 14 telomeric ends across 13 scaffolds. The final genome encompassed a total of 22,854 protein-coding genes. This is the first high-quality, haplotype resolved genome assembly of the Drakensberger breed. This assembly high accuracy, contiguity and completeness place the genome among the highest-quality cattle genomes produced to date. This genomic resource is essential in designing programs to studies for breed evolution, adaptation, genomic selection and for conservation of the South African indigenous resources.

## Introduction

The Drakensberger is a South African indigenous cattle breed which was developed over 500 years ago from cross breeding of Dutch Groningen bulls (*Bos taurus*) with Sanga-type cows (*Bos indicus*) kept by the Khoikhoi people^1^. Through careful selection, Drakensberger cattle was refined for economically important traits, including low birth weights that ease calving, strong maternal ability, moderate temperament, and natural resistance to ticks and tick-borne diseases due to their thick skin^2,3^. It is highly adaptable to sourveld grazing and harsh South African climates^4,5^. Drakensberger cattle breed is characterized by a solid black coat, long bodied muscular build, and medium-to-large frame size without a hump^6,7^. This breed has short and strong legs with hard hooves that are often compared to those of a buffalo, allowing it to walk in difficult terrain^5^. The average birth weights range from 34.8–36.3 kg for male calves and 32.8– 34.0 kg for females. At 205 days (weaning), bulls weigh between 213.5 and 232 kg, while heifers weigh 197.5–214 kg^2,5,8,9^. Mature bulls typically weigh 820–1100 kg, and cows 550–720 kg making them more popular in the beef production industry. The Drakensberger is amongst many of the African indigenous cattle breeds with good economic traits but have limited genetic information, thus prompt the need to develop a high-quality genetic reference.

The long HiFi reads genome reference at a chromosomal level are essential to inform breeding and conservation programs, especially in the livestock sector. The current available^10^ bovine genome reference was developed using a non-African breed (Hereford)^10^ which might not explain the genetic architecture of the African indigenous cattle such as the Drakensberger cattle. Some of the African indigenous cattle breeds have draft genomes which have been developed using PacBio HiFi^11,12^. These genomes might also not explain the Drakensberger cattle genome due to effects such as different agroecological regions and genetic diversity. It is essential that each breed has its own breed specific genome reference to fully leverage on the breed uniqueness. In this study, we have developed the Drakensberger genome reference using a trio-binning approach to generate a fully phased haplotypes.

## Materials and methods

The sample collection procedure including processing and handling of the animals used in this study was approved by the University of South Africa ethics committee (reference number: AREC-100818-024), Limpopo department of Agriculture and the Department of Agriculture Land Reform and Rural Development (DALRRD) under section 20 of the Animal Diseases Act 1984 (Act 35 of 84) (ref no 12/11/1/1/23 (6508 AC). Furthermore, the ethics procedure was guided by the AfricaBP policy on ethics which emphasizes ethical, legal, and social issues throughout the research activities. Finally, this work benefited from compliance consultations with the Department of Forestry, Fisheries and Environment (DFFE) as the Competent National Authority for biodiversity framework and digital sequence information in South Africa.

### Sample collection and sequencing

In this study, a pure Drakensberger breed of trio (Sire, Dam and Progeny) samples were collected (**Figure 1**) from a stud famer in Gauteng province of South Africa. Blood was collected from the tail in EDTA tubes and placed on dry ice, immediately transported to Inqaba Biotech laboratories in Pretoria, South Africa, and stored at −80°C freezer until further processing. High molecular weight genomic DNA was extracted from 200 ul blood using Nanobind protocol^13^ for whole blood high molecular weight (HMW) DNA extraction to construct a sequencing library.

**Figure 1.**
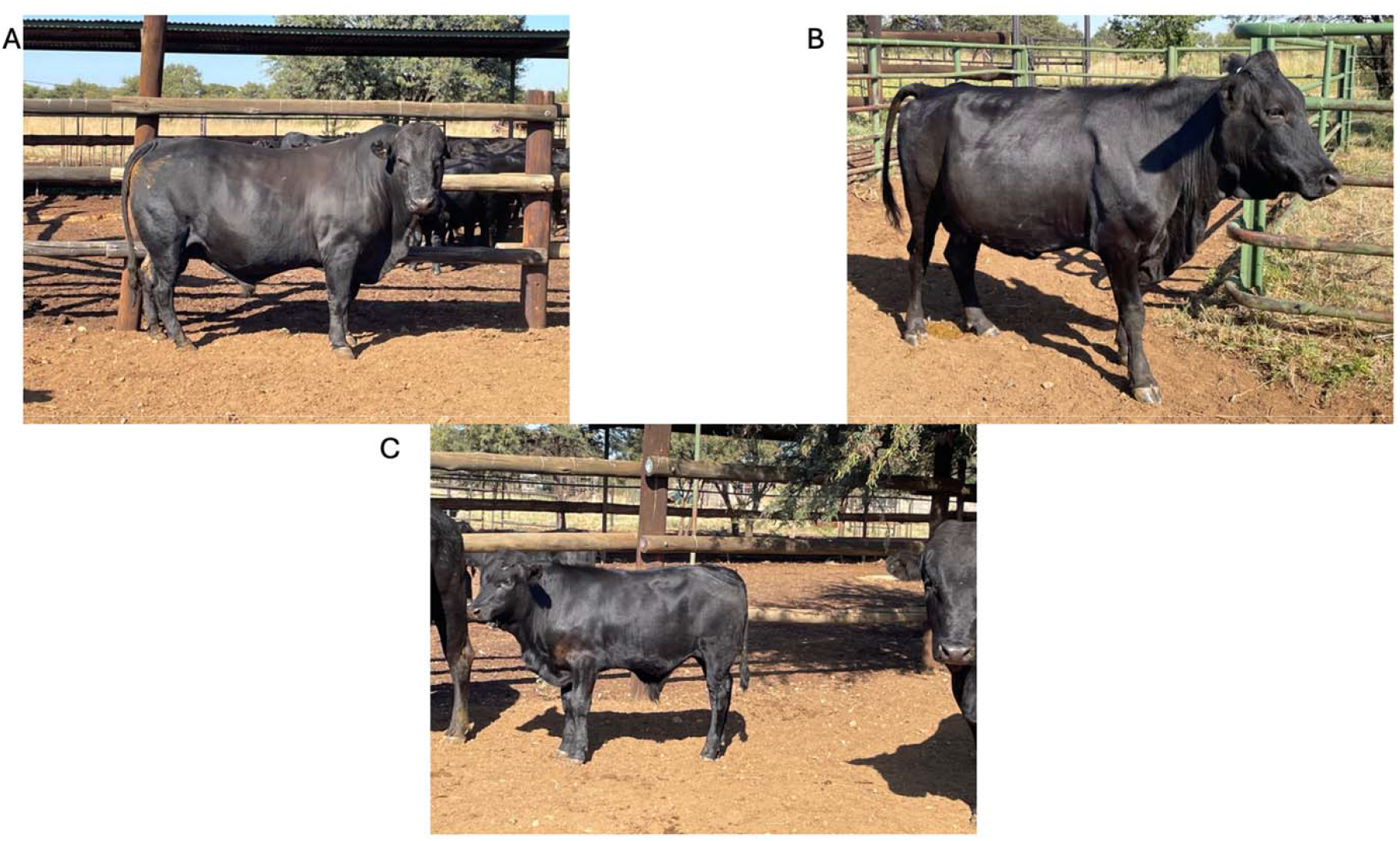
The picture of the sampled Drakensberger cattle (A=Sire, B=Dam, C= Progeny).

**T**he sire and Dam were sequenced on Pacbio revio for HiFi reads, the progeny was sequenced on PacBio revio for HiFi reads, Oxford Nanopore technologies for altra long reads and NovaSeq 6000 instrument (Illumina) to generate Omni-C reads. The protocol was optimized for extraction from 200 ul of whole blood. Library sequencing on the PacBio revio platform was done using SMRTbell® prep kit 3.0^14^ following the manufacturer’s instructions. Dovetail Omni-C library prep was performed from the same sample used for HiFi sequencing following the manufacturer’s instructions. The resulting library was sequenced on NovaSeq 6000 instrument (Illumina). For Oxford Nanopore Technologies, the sequencing was done on promethION.

### Oxford Nanopore Technologies sequencing Library preparation

DNA was first subjected to an end-prep reaction to repair ends and add ligation adapters by incubating with the NEBNext Ultra II End Repair/dA-Tailing Module for 5 minutes at room temperature followed by 5 minutes at 65°C. The reaction was purified using 60 µL of AMPure XP beads with a 5-minute binding incubation. The bead pellet was washed twice with 80% ethanol, and the DNA was eluted in 61 µL of nuclease-free water and incubated at 37°C for 10 minutes to ensure complete resuspension. For adapter ligation, the end-prepped DNA was combined with NEBNext Quick T4 DNA Ligase, Ligation Buffer, and the ONT Native Barcoding Kit. The reaction was incubated at room temperature for 10 minutes. A subsequent clean-up was performed using 40 µL of AMPure XP beads. The ligated library was then washed twice with 250 µL of Short Fragment Buffer (SFB) to remove excess adapters and eluted in 32 µL of Elution Buffer. The final library was quantified using a Qubit fluorometer and loaded directly onto PromethION flow cell for sequencing.

### Genome Assembly

The current genome assembly of the Drakensberger cattle was first carried out in the Vertebrate Genome Project (VGP) workflow^15,16^ in Galaxy platform. We first subjected the trio HiFi reads to GenomeScope^17^ for genome profiling (Figure 2) to estimate the genome size, heterozygosity and homozygosity. We then employed hifiasm^18^ on trio mode to assembly a phased genome which produced phased haplotypes (haplotype1= parental and haplotype2= maternal). To further polish the assembly, the genome was polished using ragtag^19^ then further ran through racon and medaka. Firstly, we mapped ONT reads to the final assembly which was followed by polishing with Racon for two times then polished by Medaka for the final assembly. This was further scaffolded with Omni-C reads using YaHS^20^ then ran the decontamination using dual decontamination preparation workflow which make use of Kraken2^21^. The dual decontamination workflow generates scaffolds versus Omni-C reads contact map with both haplotypes for curations. Pretextview^22^ was used to manually curate the scaffolds and by orienting and correcting mis-assemblies using the genomic proximity signal of the Omni-C reads alignment against scaffolds. The genome completeness was further assessed using BUSCO with a dataset lineage of cetartiodactyla_odb10, which had 97.5% complete genes (Figure 2).

**Figure 2.**
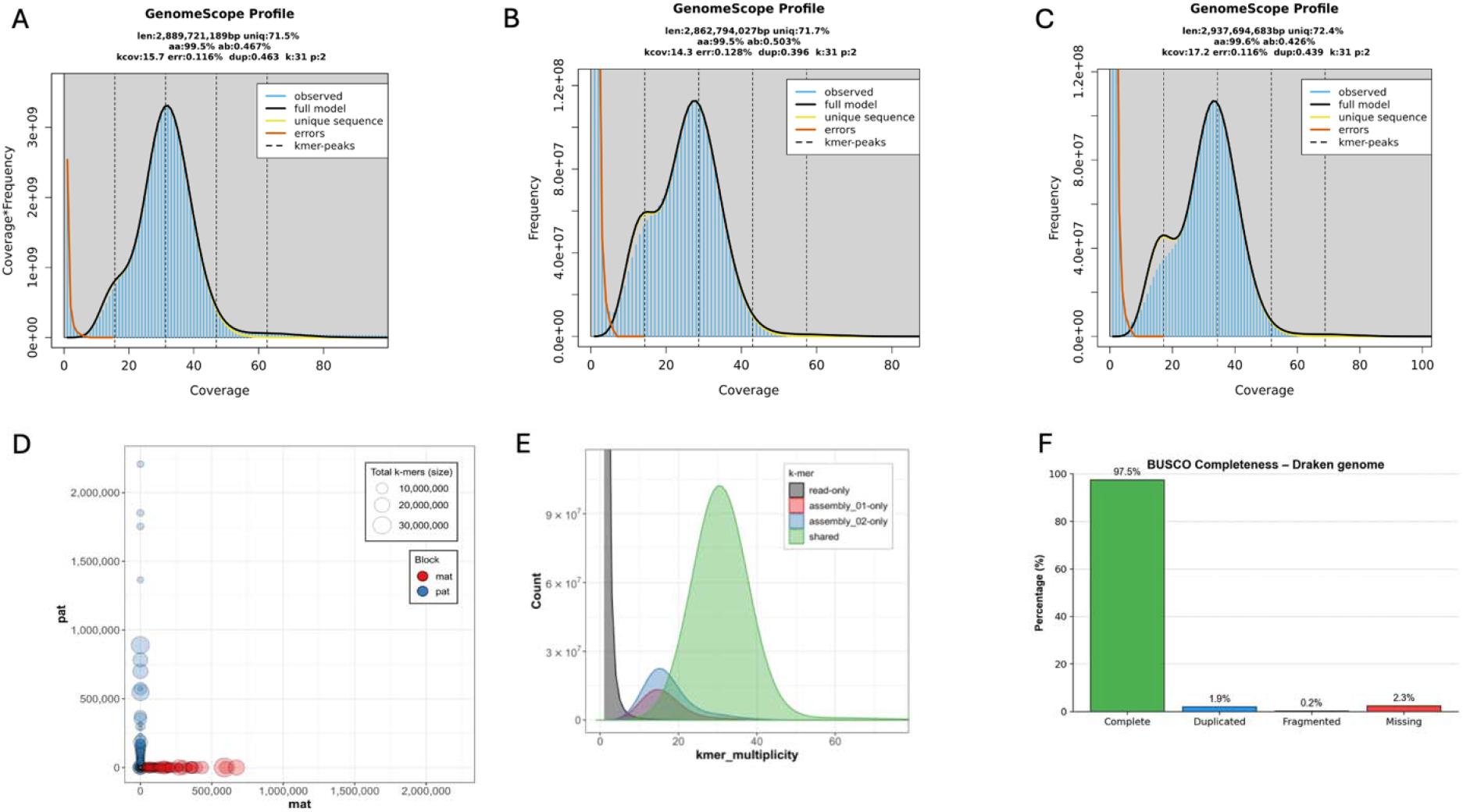
the genome profiling of the Drakensberger cattle. The main peak corresponds to the homozygous peak, used to estimate genome size for A = Progeny (∼2.89◻Gb), B = Sire (2.86 Gb) and C = Dam (2,94 Gb), and the k-mer frequency distribution for maternal and paternal (D), Progeny (E) with BUSCO completeness of the Drakensberger cattle (F).

The final genome size of the Drakensberger cattle is 2.89◻Gb. the genome size is within the expected genome size of the bovine as compared to the bovine reference genome for the Hereford^10^ which is 2.8 Gb as reported in Table 1 below. The quality of the assembly was further assessed using Merqury completeness, which was 99.5% for both assemblies for progeny and above 99% for both parents (Table 2). We further assessed the similarity of the assembled Drakensberger genome with the current bovine genome reference using D-genies v 1.5.0^23^, which uses Minimap2 v v2.28^24^. To visualize the sequence identity between the query genome and the reference genome, we used a circos plot and dot plot as reported in figure 4 below that was generated in R v 4.4.1 using the package, circilize v0.4.16^25^. The synteny plot show consensus, and the scaffolds which correspond to the reference chromosomes. Further granularity represented in the dotplot show that there are large segments which align and have similarity of about 75%. This shows the genetic variation captured in the present assembly.

**Table 1.** Assembly stats for Drakensberger cattle as compared to the assembly stats for Genome references of Hereford and Tuli cattle.

| Metrics | Drakensberger | NCBI RefSeq<br>assembly<br>(Hereford) | Progeny (Tuli x<br>Wagyu) _NCBI |
| --- | --- | --- | --- |
| Genome size | 2.89 Gb | 2.8 Gb | 3.0 Gb |
| Number of<br>scaffolds | 173 | 1,957 | 878 |
| Scaffold N50 | 111 Mb | 103.3 Mb | 101.4 Mb |
| Number of contigs | 357 | 2,343 | 909 |
| Contig N50 | 61 Mb | 26.4 Mb | 95.5 Mb |
| GC percent | 43.08% | 42% | 44% |
| Gaps | 0.001% | - | - |
| QV | 70.23 | - | - |

**Table 2.** Assembly quality results of merqury completeness.

| Assembly | k-mer<br>set | Solid k-mers in<br>assembly | Solid k-mers in<br>reads | Completeness<br>% |
| --- | --- | --- | --- | --- |
| assembly_01 | all | 2045448978 | 2349517914 | 87.0582 |
| assembly_02 | all | 2172481400 | 2349517914 | 92.465 |
| both | all | 2336864728 | 2349517914 | 99.4615 |
| assembly_01 | pat | 75061950 | 77333815 | 97.0623 |
| assembly_01 | mat | 207127 | 25052032 | 0.826787 |
| assembly_02 | pat | 22025085 | 77333815 | 28.4805 |
| assembly_02 | mat | 24806096 | 25052032 | 99.0183 |
| both | pat | 76677538 | 77333815 | 99.1514 |
| both | mat | 24961346 | 25052032 | 99.638 |

### Genome annotation of Drakensberger cattle

The genome was annotated using a deep learning-based ab initio gene structure prediction tool, Tiberius^26^, which generated gene predictions based on inherent structural evidence. This resulted in 22,854 predicted protein-coding genes for the Drakensberger as compared to the 25,591, 24,367 and 64,745 for the Nguni, Bonsmara and Hereford cattle respectively. The distribution and unique orthogroups found in the Drakensberger as compared to the Hereford, Bonsmara, and Nguni were reported in figure 5. A total of 15,755 orthogroups are common to all breeds, with each breed also exhibiting distinct sets of unique orthogroups, reflecting both shared genomic architecture and breed-specific orthogroups. Furthermore, we assessed the abundance of the structural variants (SV) detected in the Drakensberger cattle as compared to the Bovine reference, Hereford cattle^10^ as reported in figure 6. In the SV, the Drakensberger cattle contains more insertions than deletions, this might suggest that Drakensberger has new or unique genomic sequences as compared to the reference genome. The insertions in the Drakensberger might be breed specific adaptations related to traits such as environmental resilience to heat stress, diseases resistance and fertility. The categories of repetitive elements identified in the Drakensberger cattle genome, and a distribution of repetitive elements are summarized in figure 7.

**Figure 4.**
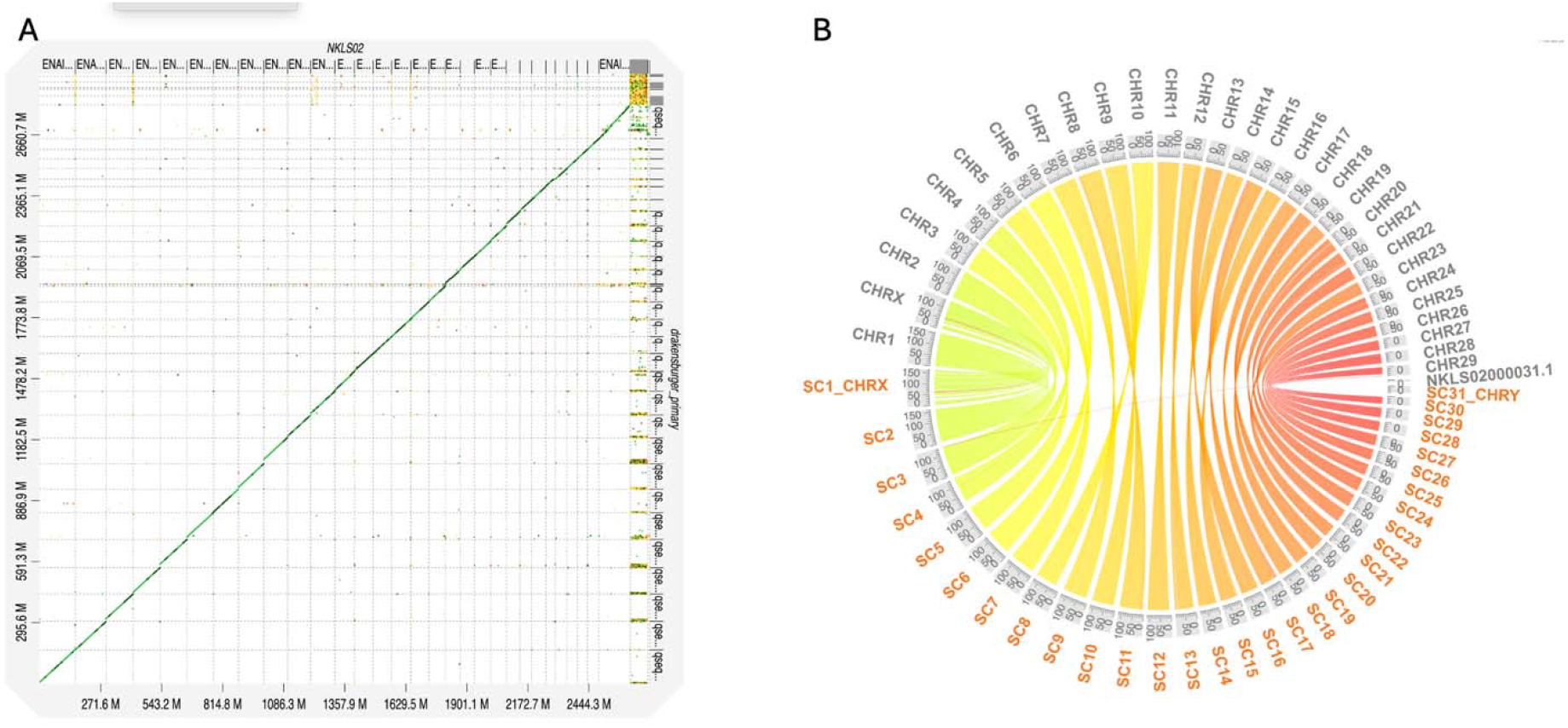
Dot-Plot (A) showing sequence identity between the Drakensberger and the reference assembly ARS2.0 (Hereford). (B) Circos plot showing comparison of synteny between the Drakensberger cattle and the reference assembly ARS2.0 (Hereford)

**Figure 5.**
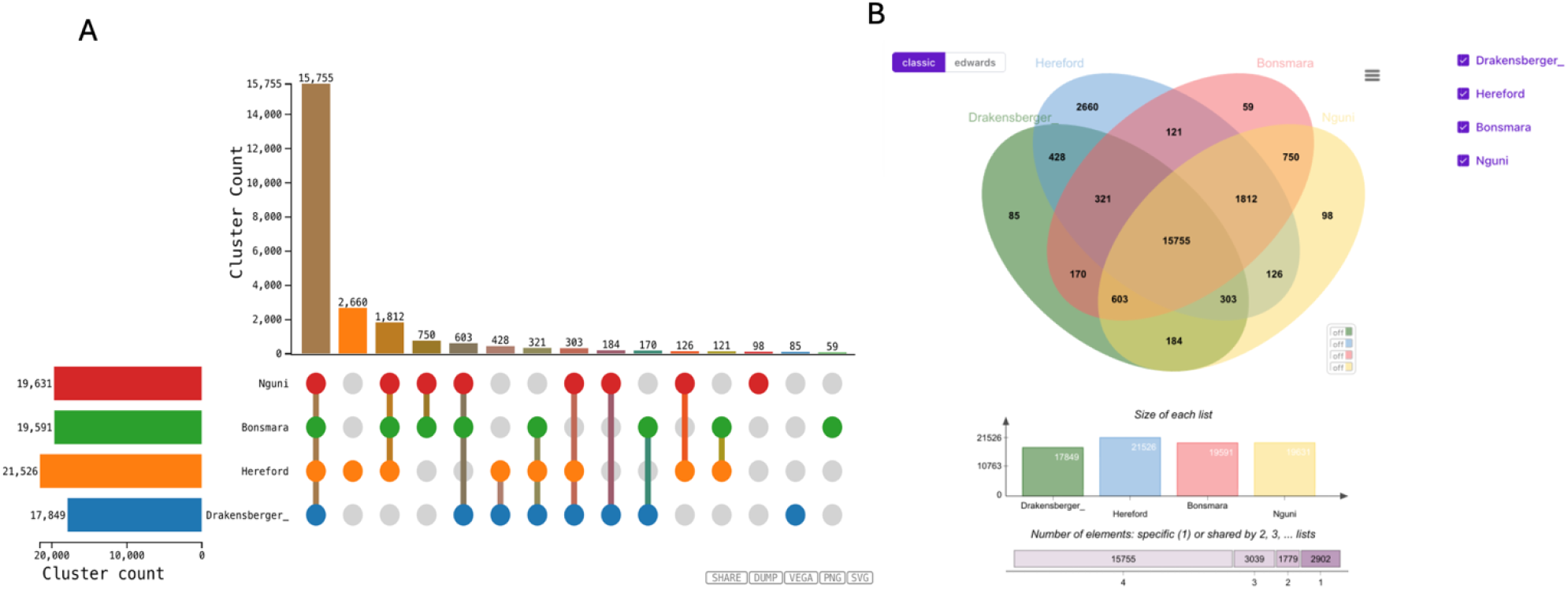
Distribution of Shared and Breed-Specific orthogroups in Drakensberger, Hereford, Bonsmara, and Nguni. The UpSet plot (A) quantifies unique and shared orthogroups clusters across different breed combinations, while the Venn diagram (B) visually summarizes their intersections.

**Figure 6.**
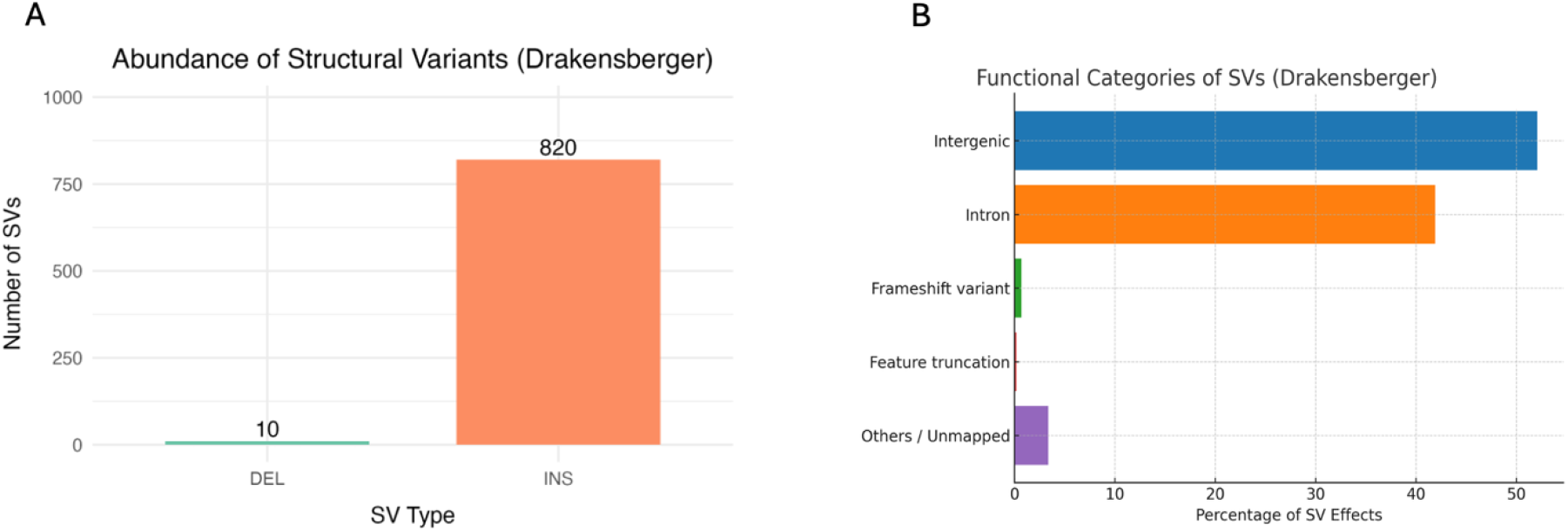
Structural Variant Landscape in the Drakensberger Population. Abundance (A) and Functional categories of Structural variants of Drakensberger cattle (B).

**Figure 7.**
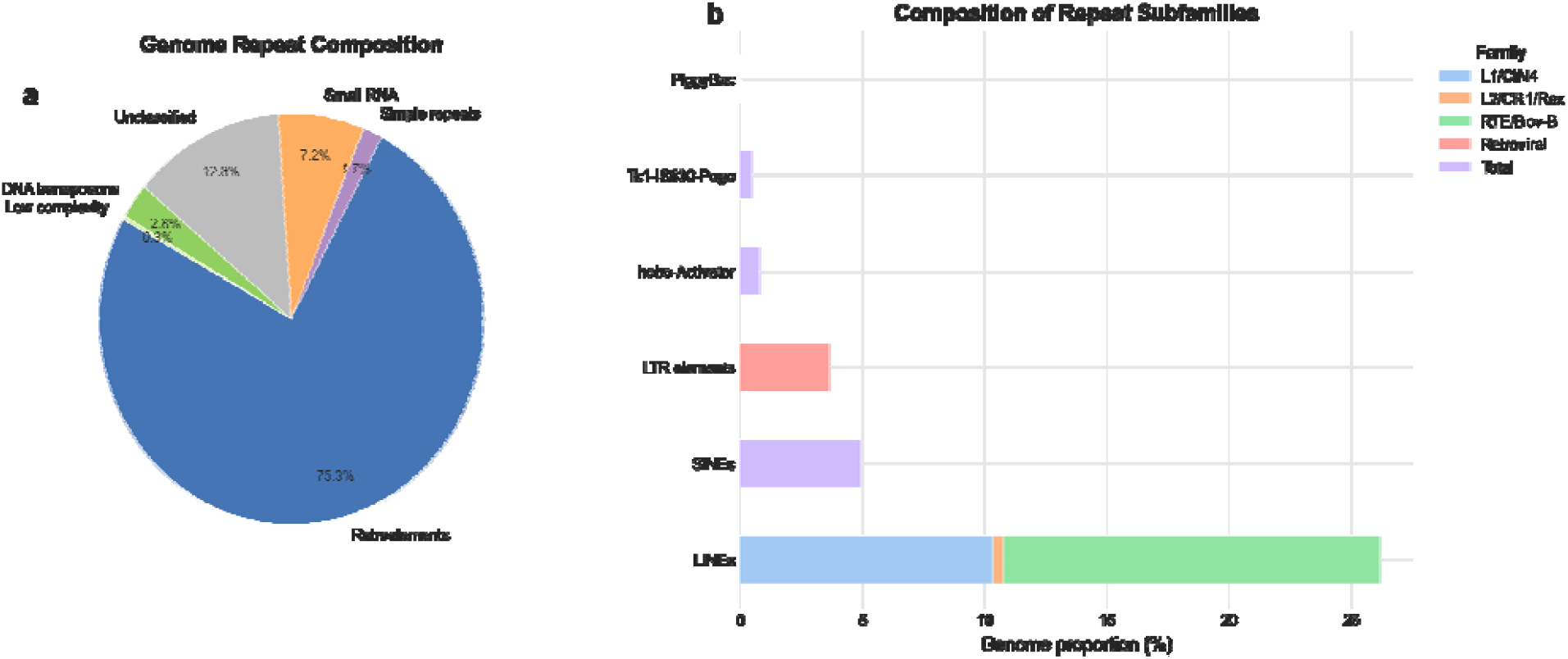
Repetitive element distribution in percentage for the Drakensberger cattle. Genome repeat composition (a) and composition of repeat subfamilies (b).

This is the first high-quality chromosomal level genome assembly of the Drakensberger cattle. The assembly stats generated in this genome is among the top assemblies and comparable if not better so the current available cattle genome reference. Our research provides a near complete telomere-to-telomere reference genome which provides potentials to advance genetic studies and breeding programs for Drakensberger cattle and as reference to other cattle breeds.

### Data Records

The raw sequences were submitted to the NCBI under the submission number: SUB1577423. The final genome assembly for the Drakensberger cattle was submitted to the NCBI with the Project number: SUB15768644.

### Technical Validation

The reference genome assembly of Drakensberger cattle was supported by VGP workflows for genome assembly which were designed to reduce human error by employing workflows that produces near error-free high-quality reference genome. The assembly was done using Hifiasm on a trio mode. The pipeline is comprised of different workflows that make use of methods such as BUSCO and mercury to assess the integrity of the genome.

## Code availability

All the analyses done in the current study were processed by employing the VGP workflows that are publicly available in galaxy (https://galaxyproject.org/projects/vgp/workflows/). All the commands and pipelines were executed following the manual and protocols of the corresponding bioinformatics software. In all the workflows, unless mentioned and where necessary, we used the default parameters.

## Author contribution

NM = Designed, sourced funding, wrote the main manuscript. TT= did the annotation and sequenced the genome. RS= prepared figures. NH= generated structural variance data. LTN= wrote the main manuscript, prepared figures, supervised the data collection and did the genome assembly.

